# Some like it hotter: differential thermal preferences among lizard color morphs

**DOI:** 10.1101/2022.11.11.516149

**Authors:** Asher Thompson, Vassiliki Kapsanaki, Heather E. M. Liwanag, Panayiotis Pafilis, Ian J. Wang, Kinsey M. Brock

## Abstract

Temperature rules the lives of ectotherms. To perform basic biological functions, ectotherms must make behavioral adjustments to keep their body temperatures near a preferred temperature (Tpref). Many color polymorphic lizards are active thermoregulators and exhibit morph differences in traits related to thermoregulation, such as color, body size, and microhabitat use. The Aegean wall lizard, *Podarcis erhardii*, is a heliothermic lizard with orange, white, and yellow color morphs that differ in size, behavior, and microhabitat use. Here, we tested whether *P. erhardii* color morphs from the same population from Naxos island, Greece, differ in Tpref. We hypothesized that orange morphs would prefer lower temperatures than white and yellow morphs because orange morphs are often found on cooler substrates and in microhabitats with more vegetation cover. We obtained Tpref for 95 individuals using laboratory thermal gradient experiments of wild-caught lizards and found that orange morphs do, indeed, prefer significantly cooler temperatures, regardless of body size differences. Average orange morph Tpref was 2.3 ºC lower than average white and yellow morph Tpref. Our results add support to the idea that *P. erhardii* color morphs have multivariate alternative phenotypes and present the possibility that thermally heterogeneous environments play a role in the maintenance of color polymorphism in this species.

## INTRODUCTION

More than 98% of all animal species are ectothermic and have strong biological ties to temperature (Angilletta et al., 2002; Angilletta, 2009). To maintain an appropriate body temperature, many ectothermic reptiles thermoregulate by basking in the sun, sheltering in shade, and moving between warmer or colder areas (Pough & Gans, 1982). For reptiles, basic biological processes such as growth, digestion, egg production, locomotion, and mating and escape behavior are all affected by body temperature (Huey, 1982; Cooper, 2000). Thus, body temperature is arguably the most important ecophysiological variable for reptiles (Stapley, 2006; Meiri et al., 2013; Garcia-Porta et al., 2019).

Thermal preference represents an important parameter of ectotherms’ thermal profiles and has been interpreted as an intrinsic character (Van Damme et al., 1986; Carneiro et al., 2017). Since many of its bodily functions and behaviors are temperature-sensitive, knowing an ectotherm’s preferred body temperature is key to understanding many aspects of its life (Taylor et al., 2020). Tpref is also considered a significant ecological index that may be linked to species distributions (Buckley et al., 2010; Crickenberger et al., 2020). In nature, an animal’s available range of temperatures is governed by its access to the diversity of microhabitats available to it, including cooler areas near water or in shade or hotter areas in full sun. Many ectothermic animals, such as insects, snakes, toads, and lizards, exhibit individual variation in their preferred temperatures and microhabitat selection based on their morphologies, coloration, and social behavior (Forsman, 2000; Bittner et al., 2002; Ng et al., 2013; Sanabria et al., 2014). Intraspecific color morphs with morph-associated morphology, social behaviors, and microhabitat associations have evolved in many species across the tree of life (Hugall & Stuart-Fox, 2012; Brock et al., 2022c), particularly in lizards (Stuart-Fox et al., 2020). Yet, relatively little is known about intraspecific color morph variation in preferred body temperature (but see Paranjpe et al., 2013; George & Miles, 2022). Indeed, animal color morphs are expected to use different subsets of available resources and occupy, on average, different environmental niches due to morph-associated morphological and microhabitat differences (Forsman et al., 2008). Color polymorphic lizards provide an opportunity to understand the relationships among color, microhabitat selection, and thermal preference.

Color polymorphism is the evolution of two or more genetically-determined color morphs within a single population (Ford, 1945; Forsman et al., 2008). Lizards from distantly related families have independently evolved similar color morphs that usually have orange, yellow, blue, or white throats (Stuart-Fox et al., 2020; Brock et al., 2022c). Color morphs often differ in more than just color; they may also exhibit differences in other traits, including body size, behavioral and physiological differences, and microhabitat preferences (Gray & McKinnon 2007; Stuart-Fox et al., 2020). When morphs with co-adapted trait complexes, or alternative phenotypes, coexist in a population, they may seek out different microhabitats with different visual properties to maintain crypsis from visual predators or to maximize visual signal efficacy to conspecifics (Leal & Fleishman, 2004; Bond & Kamil, 2006; Maan et al., 2008; Ng et al., 2013). For example, in *Anolis* lizards, signal efficacy for brighter dewlap colors like reds, pinks, and oranges increases in darker, mesic habitats, which are also usually cooler than sunnier, open xeric habitats (Leal & Fleishman, 2004; Ng et al., 2013). If color morphs differ in the brightness of their color patch and occupy distinct microhabitats with different thermal conditions, then morphs may have different temperature preferences and thus exhibit other life history differences.

The Aegean wall lizard, *Podarcis erhardii*, is a color polymorphic lacertid lizard that behaviorally thermoregulates to maintain its body temperature (Belasen et al., 2017; Pafilis et al., 2019; Brock et al., 2020). Like many *Podarcis* lizards, adult *P. erhardii* have three genetically determined throat color morphs that are orange, white, or yellow (Andrade et al., 2019; Brock et al., 2022c; Figure 1). *Podarcis erhardii* color morphs differ in throat color brightness (Brock et al., 2020), body size, behavior (Brock et al., 2022a), and habitat use (BeVier et al., 2022; Brock & Madden, 2022), which are traits that could have implications for morph differences in thermal preferences (Digby, 1955; Forsman, 2000). In general, *P. erhardii* orange morphs and white morphs tend to be most different from each other in these traits, and yellow morphs usually exhibit intermediate phenotypes in a suite of physical and behavioral traits (Brock et al., 2020). Orange males are larger than white and yellow males, and female color morphs do not differ significantly in size (Brock et al., 2020), though orange females tend to be on the larger end of the size spectrum. Orange morphs are found more often in microhabitats that have more vegetation and are cooler; in contrast, white and yellow morphs are more often found in areas that are open and hotter (BeVier et al., 2022). In laboratory behavioral experiments conducted on male color morphs, orange morphs were the least bold and aggressive, yellow morphs were intermediate, and white morphs were the most active, aggressive, and adept at winning contests over limited open basking space (Brock et al., 2022a). These differences in *P. erhardii* color morph morphology, microhabitat selection, and social behavior may play a role in their thermoregulatory behavior and thermal preferences. Variation in color morph thermal preferences may have profound implications for color morph life histories and relative color morph fitness in hotter, drier environments.

**Figure 1.**
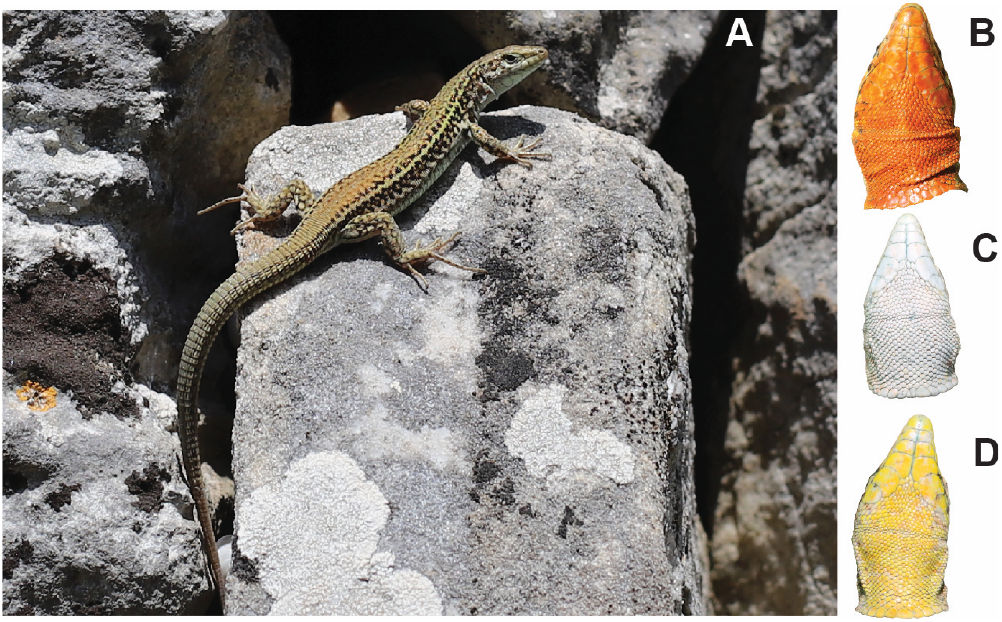
Study species. A) A white morph *Podarcis erhardii* basking on a dry stone wall, Naxos, Greece. Images of *P. erhardii* orange (B), white (C), and yellow (D) throat color morphs.

Here, we quantify color morph Tpref with thermal gradient experiments in the lab to determine if *P. erhardii* color morphs exhibit different preferred temperatures. We hypothesize that orange color morphs prefer cooler temperatures than white and yellow morphs due to their more frequent use of darker, cooler microhabitat. Given their similar choices for microhabitat that is open and hotter (Brock & Madden, 2022; BeVier et al., 2022), we hypothesize that white and yellow morphs would not have significantly different preferred temperatures from one another. Past experimental and natural observations of *P. erhardii* behavior indicate that white morphs are more active than other morphs (Brock & Madden, 2022; Brock et al., 2022a); because of this, we also hypothesized that white morphs should shuttle more between cooler and hotter temperatures.

## METHODS

### Study Species

*Podarcis erhardii* is a small to medium-sized (adult SVL range: 45-75 mm) lacertid lizard (Figure 1) that is highly variable in color and body size across its discontinuous range comprising hundreds of islands in the Aegean Sea (Marshall et al., 2015; Stadler et al. 2022, in press; Brock et al., 2022b). Adults have one of six genetically-determined throat colors: orange, orange-white, white, white-yellow, yellow, or yellow-orange (Andrade et al., 2019; Brock et al., 2020). We limited our study to the three monochromatic orange, white, and yellow morphs to maximize genetic and phenotypic differences. This lizard is diurnal and has an activity period from 08:00 to 19:00 in summer, when the days are longest and hottest (Belasen et al., 2017). As its vernacular name suggests, the Aegean wall lizard is commonly found basking on dry stone walls that are common throughout the region.

*Podarcis erhardii* is endemic to the southern Balkans and is present on many of the Aegean islands, including the Cyclades and Sporades island clusters. This region experiences a classic Mediterranean climate with a long dry period that commences at the end of spring, hot summers, and cool wet winters (Valakos et al., 2008). *Podarcis erhardii* is a generalist that can be found in a variety of habitats, from low-elevation sandy dunes with sparse vegetation, to mid-elevation rocky, dry areas with dense xeric vegetation, all the way up to forested montane habitats reaching 1400 meters in elevation (Lymberakis et al., 2018).

### Study Area and Sampling

In May 2022, we sampled 95 lizards from Naxos island, Greece. Naxos is the largest island in the Cyclades island cluster (land area: 440 km^2^) and is the center of *P. erhardii’s* distribution in the Aegean Sea. We sampled lizards from the terraced foothills below Profitis Ilias peak (elev. 590 m a.s.l., 37.08043203432896 ºN, 25.49171180549804 ºE).We chose this site because it features a diverse combination of habitat types that represent the range of thermal microhabitats available throughout the island. The site is located around a remote hiking path bordered by dry stone walls, which also form the terraced agricultural plots at the site. The vegetation at this site consists of a mixed matrix of grasses, phrygana (*Euphorbia acanthothamnos*), olive trees (*Olea europaea*), and sclerophyllous evergreen maquis. This diversity of vegetation, combined with stone walls and hiking paths that wind through hot open areas, provides a range of temperatures for *P. erhardii* to behaviorally thermoregulate.

We sampled only adult lizards (> 45 mm SVL) for this study. Lizards were caught with a lasso attached to the end of a telescopic fishing pole. Color morph is discernible by eye (Brock et al., 2020), and we categorized morphs as orange, white, or yellow in the field upon capture. Prior to experimentation, we measured lizard snout-vent length (SVL) in mm with a pair of digital calipers (Mitutoyo 500-171-30 Absolute Scale Digital Caliper, Aurora, Illinois, USA) and recorded their mass to the nearest 0.1 g with a digital scale (Pesola PTS3000 Platform Scale, Schindellegi, Switzerland). Female lacertid lizards lack active femoral pores, and so we sexed lizards based on the absence or presence of femoral pores and hemipenes. After preferred temperature experiments, we returned animals to their exact point of capture. The University of California, Berkeley IACUC (protocol AUP-2021-08-14567) and the Greek Ministry of Energy and the Environment (permit YΠEN/ΔΔΔ/5619/145) approved animal handling and use protocols used in this study.

### Preferred Temperature Experiments

We conducted preferred temperature experiments in an air-conditioned room at 19 ºC with constant lighting between 8:00 and 20:00. To measure Tpref, we used a specially-designed thermal gradient made of cardboard measuring 100 × 44 × 28 cm (Figure 2). We divided the thermal gradient into four, 11 cm wide lanes, which allowed us to test four lizards simultaneously. Lanes were divided by opaque cardboard so that lizards could not see each other and to reduce the potential effects of visual cues on positioning and thermoregulatory behavior. We positioned heating bulbs in order from left to right of 65 W, 150 W, and 65 W directly over each of the 3 lane dividers at one end of the gradient (Figure 2). We placed ice packs at the opposite end to create a thermal gradient from 15 ºC to 45 ºC and confirmed gradient temperatures (measured by thermometer, see Van Damme et al., 1986). We covered the floor of the temperature gradient in one centimeter of sand from the island to give lizards traction for free movement and a more natural and familiar substrate. We replaced the sand between every experiment to remove any waste or scents from the previous lizard that might affect thermoregulatory behavior. We did not feed or provide water to lizards prior to experimentation.

**Figure 2.**
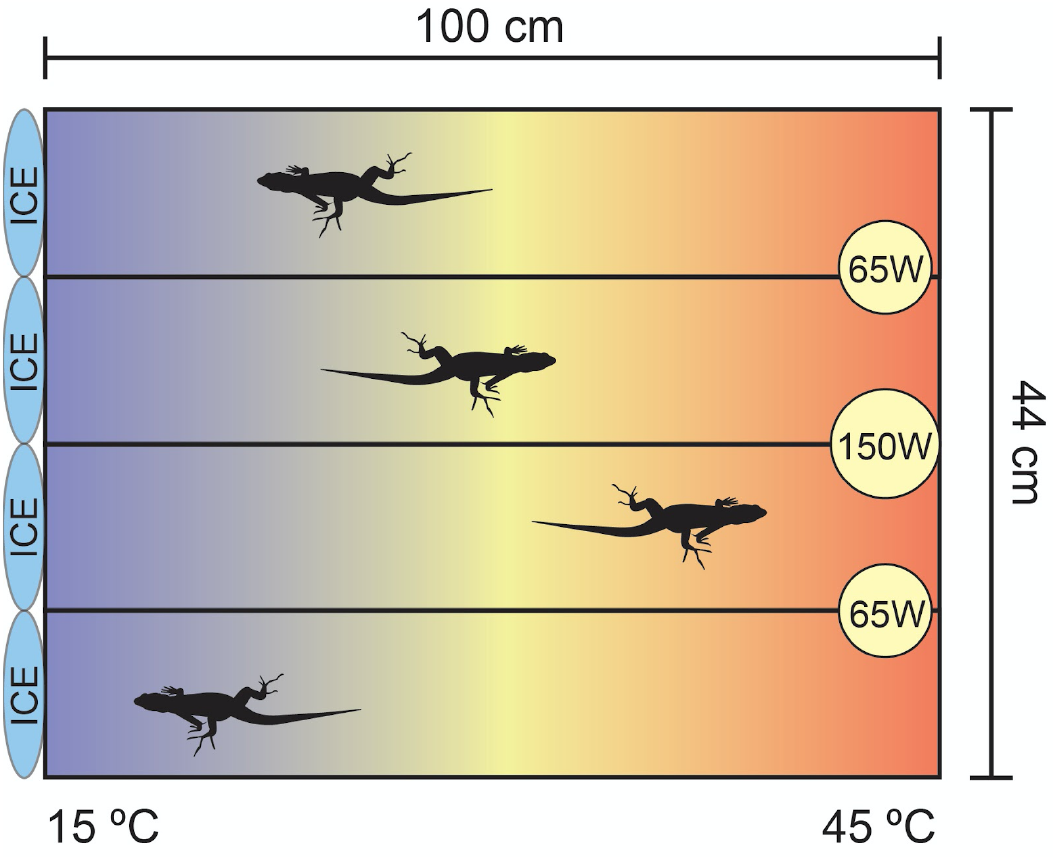
Experimental set-up. Diagram of the thermal gradient used to measure Tpref in *P. erhardii*, as viewed from above. The gradient temperature ranged from 15 ºC at the cold end to 45 ºC at the hot end. The gradient was maintained with ice packs (blue ovals) at the cold end, and heating bulbs of 65 W and 150 W (yellow circles) at the hot end.

To begin a Tpref experiment, we placed a lizard in the middle of a lane and allowed it to acclimate to the thermal gradient for 30 min (N = 52) without taking any body temperature measurements (Carretero, 2012; Carneiro et al., 2015; Belasen et al., 2017). For a subset of experiments we extended the acclimation period to 60 min (N = 19) and 120 min (N = 24) to determine if acclimation period significantly affected Tpref. We randomized lane assignments and ran experiments with sets of individuals that varied by morph, sex, and reproductive status. After the acclimation period, we took external body temperature measurements with a FLIR TG5 temperature gun (Hare et al., 2007; Trochet et al., 2018; George & Miles, 2022) every 5 min for 1 h (N = 50), 3 h (N = 24), and 6 h (N = 21). We ran trials of different lengths due to time constraints in the field and tested if individual Tpref differed between the 60 min timepoint and final timepoint in experiments lasting 3 h and 6 h. To keep body temperature measurements consistent among individuals, we took temperature readings on the dorsal surface between the lizard’s shoulders. In preliminary experiments (N = 20), a quick-reading thermocouple inserted 5 mm into each lizard’s cloaca and taped into place altered their ability to freely move freely through the gradient and their general behavior (e.g., biting the wire, inspecting the wire, and attempting to remove the tape). If a lizard moved while we took a surface temperature reading, we marked that measurement N/A. We used each lizard in only one Tpref experiment.

We calculated individual Tpref for each lizard as the 50% interquartile range (IQR) of all body temperature measurements taken while they were in the thermal gradient after the acclimation period. Color morph Tpref was calculated as the 50% IQR of all individual Tpref calculations for each color morph. We calculated a shuttling variable by summing the absolute values of the change in body temperature at 5-min intervals (George & Miles, 2022). This measurement describes the thermoregulatory behavior of individuals during the experiment. A high shuttling value indicates that individuals moved around more and basked in a wider range of temperatures throughout the gradient, and a low shuttling value indicates that individuals moved less and basked in a narrower range of temperatures throughout the gradient.

### Statistical Analyses

We performed all statistical analyses in R v4.1.1 (R core team, 2021). We checked that Tpref data met ANOVA, ANCOVA, Kruskal-Wallis, Pearson correlation, t-test, and Wilcoxon Rank Sum test assumptions using QQ plots, Shapiro-Wilk tests, and Levene’s tests. We first evaluated whether individuals exhibited any significant differences in Tpref related to sex or reproductive status. We checked for a difference in Tpref between gravid (N = 21) and non-gravid (N = 17) females with a t-test and whether Tpref differed by reproductive status for all lizards (gravid female, non-gravid female, male) using a Kruskal-Wallis test. We determined whether females and males differed in Tpref using a Wilcoxon Rank Sum test.

We then examined whether experimental conditions had any effect on observed Tpref. To determine whether differences in acclimation time in the thermal gradient had a significant effect on individual variation in Tpref, we ran a linear model with acclimation time and experiment length as explanatory variables. We also tested whether Tpref differed between experiments of different lengths (1 h, 3 h, and 6 h) for each morph using one-way ANOVA and Kruskal-Wallis tests. For experiments lasting 3 h and 6 h, we tested for individual differences in average Tpref at 1hr and at the end of the trial with paired t-tests. We used Pearson correlation tests to test for significant correlations between Tpref and the time of the experiment and between Tpref and the length of time between capture and experiment.

Lastly, we checked whether two additional factors, position in the experimental chamber and body size, had any effect on Tpref. To ensure there were no significant effects of the different gradient lanes, we tested for an effect of the gradient lane on Tpref using a linear model with lane coded as a factor. Finally, because body size is often associated with thermal physiology in lizards, we tested for correlations between body size metrics (mass and SVL) and Tpref using Pearson correlation tests.

To test whether Tpref differed significantly by color morph, we ran a one-way ANOVA followed by a *post-hoc* Tukey HSD test. We also ran ANCOVA tests with mass and SVL as covariates to control for differences in body size in morph comparisons of Tpref.

For our shuttling variable data, we checked that these data met Wilcoxon Rank Sum and Kruskal-Wallis test assumptions using Shapiro-Wilk tests and Levene’s tests and removed outliers when necessary to meet those assumptions. We investigated sex differences in shuttling behavior using a Wilcoxon Rank Sum test (N = 36 females, N = 54 males). To determine whether color morphs differed in their shuttling behavior, we used Kruskal-Wallis tests on data from the first 60 min of all experiments (N = 90).

## RESULTS

We obtained Tpref estimates for 95 lizards. We removed six outliers that were more than two standard deviations away from the mean to achieve normality and meet ANOVA, ANCOVA, Kruskal-Wallis, Pearson correlation, t-test, and Wilcoxon Rank Sum test assumptions, resulting in a final dataset with 89 lizards (Table 1). Tpref ranged from 29.7 ºC to 37.8 ºC (mean = 34.8 ± 1.85 ºC). Tpref did not significantly differ between gravid (Tpref = 35.2 ± 1.9 ºC) and non-gravid (Tpref = 34.6 ± 2.0 ºC) females (Welch’s two sample t-test, *t* = -0.899, df = 33.8, *P* = 0.375), reproductive status of all lizards (Kruskal-Wallis test, *χ2* = 1.42, df = 2, *P* = 0.49), females and males (Wilcoxon Rank Sum test, *W* = 1039.5, *P* = 0.561), nor experiments of different lengths for each morph (one-way ANOVA, orange: *F* = 2.91, df = 1, *P* = 0.122; Kruskal-Wallis test, white: *χ2* = 3.02, df = 2, *P* = 0.22; Kruskal-Wallis test, yellow: *χ2* = 1.06, df = 2, *P* = 0.588). We also did not detect any significant differences in Tpref measured at 1 h and 3 h or 1 h and 6 h in experiments lasting 3 h and 6 h (paired t-test, 1 h and 3 h experiments: *t* = 0.767, df = 23, *P* = 0.45; paired t-test 1 h and 6 h experiments: *t* = 0.716, df = 20, *P* = 0.48). Finally, acclimation time and experiment time were not significant predictors of Tpref (R^2^adj = - 0.009, *F* = 0.589, df = 86, *P* = 0.56). Given these results, we pooled individuals regardless of reproductive status, sex, experiment length, and acclimation and experiment time in further comparisons of color morph Tpref.

**Table 1.** Color morph, sex, sample size (n), mass, SVL, and Tpref for P. erhardii used in this study. Numerical values are presented as mean ± SD.

| Morph (n) | Mass (g) | SVL (mm) | $T_{pref}$ ($^{\circ}\text{C}$) |
| --- | --- | --- | --- |
| **Orange (29)** | $7.97 \pm 1.51$ | $65.9 \pm 3.48$ | $33.2 \pm 1.45$ |
| Females (10) | $6.66 \pm 0.636$ | $64.6 \pm 2.30$ | $32.7 \pm 1.72$ |
| Males (19) | $8.66 \pm 1.38$ | $66.7 \pm 3.83$ | $33.4 \pm 1.27$ |
| **White (30)** | $6.29 \pm 1.45$ | $61.6 \pm 4.29$ | $35.6 \pm 1.63$ |
| Females (13) | $5.37 \pm 1.26$ | $60.4 \pm 4.84$ | $35.7 \pm 1.60$ |
| Males (17) | $6.99 \pm 1.18$ | $62.6 \pm 3.69$ | $35.5 \pm 1.69$ |
| **Yellow (30)** | $6.48 \pm 1.71$ | $62.9 \pm 5.13$ | $35.7 \pm 1.29$ |
| Females (15) | $5.13 \pm 1.33$ | $59.0 \pm 5.58$ | $35.7 \pm 1.13$ |
| Males (15) | $7.83 \pm 0.69$ | $65.3 \pm 1.33$ | $35.7 \pm 1.47$ |

We found no significant correlation between individual Tpref and experiment time (Pearson’s correlation, *t* = 1.033, df = 87, *P* = 0.30) or the amount of time between capture and experiment (Pearson’s correlation, *t* = 1.633, df = 87, *P* = 0.11). Additionally, we did not detect a lane effect from the thermal gradient on Tpref (R^2^adj = -0.027, *F* = 0.235, df = 85, *P* = 0.87). Tpref was significantly negatively correlated with lizard body size (Pearson’s correlation, *t* = -3.279, df = 87, *P* = 0.001) and SVL (Pearson’s correlation, *t* = -3.386, df = 87, *P* = 0.001).

We found that preferred temperature (Tpref) differed significantly among color morphs (one-way ANOVA, F = 27.64, df = 2, *P* < 0.001, Figure 3). A post-hoc Tukey HSD test revealed that the Tpref of orange morphs was significantly cooler than white (orange-white difference = -2.39 ºC, *P* < 0.001) and yellow (orange-yellow difference = -2.52 ºC, *P* < 0.001) morphs. White and yellow morph Tpref did not differ significantly (white-yellow difference = - 0.12 ºC, *P* = 0.94). The statistically significant difference in morph Tpref remained even when controlling for mass (ANCOVA morph *F* = 20.11, df = 2, *P* < 0.001; mass *F* = 0.65, df = 1, *P* = 0.42) and SVL (ANCOVA morph *F* = 20.20, df = 2, *P* < 0.001; svl *F* = 1.39, df = 1, *P* = 0.24).

**Figure 3.**
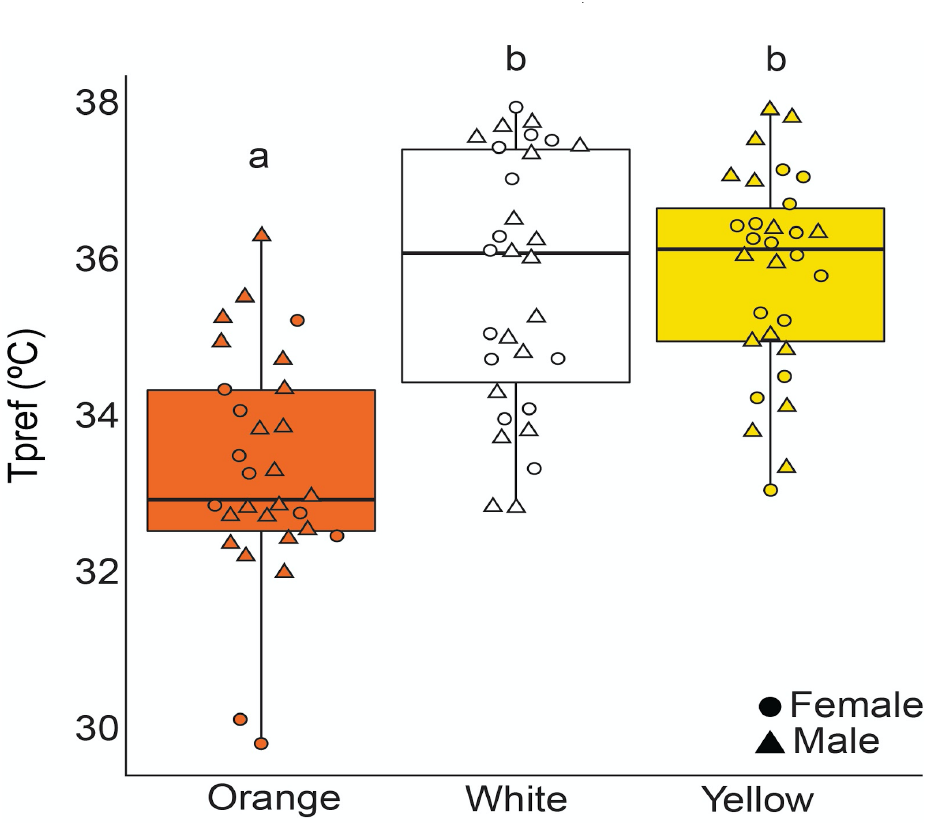
Color morph Tpref. Thermal preference (Tpref, ºC) for *P. erhardii* according to throat color morph (orange, white, yellow). Box plots show 25% and 75% quartiles (boxes), medians (lines in the boxes), and outermost values within the range of 1.5 times the respective quartiles (whiskers). Values for individual lizards are shown as circles (females) and triangles (males). Significantly different group means are indicated by different letters.

We did not find significant differences in shuttling behavior between the sexes (Wilcoxon Rank Sum test, *W* = 610, *P* = 0.776). On average, females changed their body temperature by 27.7 ± 15.2 ºC in the first 60 min of Tpref experiments, whereas males changed their body temperature by 28.1 ± 12.8 ºC during the first 60 min of Tpref experiments. We also did not find significant differences in shuttling behavior between color morphs (Kruskal-Wallis test yellow: *χ2* = 2.40, df = 2, *P* = 0.301). On average, orange morphs changed 26.5 ± 13.6 ºC in body temperature, white morphs changed 29.0 ± 9.6 ºC in body temperature, and yellow morphs changed 28.5 ± 16.6 ºC in body temperature during the first 60 min of Tpref experiments (N = 90).

## DISCUSSION

We found significant support for our hypothesis that the brighter, larger orange morphs would prefer cooler temperatures compared to the white and yellow morphs (Figure 3), which we had based on their observed preference for microhabitats that offer cooler, shadier conditions (BeVier et al., 2022). A hypothesized consequence of color polymorphism is the evolution of morphs that have non-overlapping environmental resource use and distinct thermal physiologies (West-Eberhard, 1986; Galeotti & Rubolini, 2004; Forsman & Åberg, 2008; Forsman et al., 2008). This can be due to a combination of different morph-specific traits, including differences in morph chroma and brightness that determine relative crypsis and visual conspicuousness in different lighting environments, microhabitat selection based on different heating and cooling rates of color morphs, and body size differences (Forsman, 2000; Forsman et al., 2008). In Aegean wall lizards, morph colors are restricted to the throat, which they display to other lizards by lifting their head back (Brock et al., 2022a). It is unclear what information these throat color signals contain and how much, if any, effect color variation in this body region has on heating and cooling rates. Previous work in reptiles has shown that ventral pigmentation has significant thermal effects on heating and cooling (Tanaka, 2007; Smith et al., 2016), and that differences in substrate thermal properties can influence ventral color evolution (Goldenberg et al., 2021). Throat color polymorphism has been associated with numerous trait deviations among the different color morphs in *Podarcis* lizards (Calsbeek et al., 2010; Abalos et al., 2016; Brock et al., 2020; Sreelatha et al., 2021), though its significance on thermoregulation remains understudied. Color morph differences in Tpref may have implications for long-term morph coexistence within and among populations - a phenomenon that has intrigued and puzzled biologists for decades (Ford, 1945; West-Eberhard, 1986; Gray & McKinnon, 2007).

Contrary to our expectation, color morphs did not exhibit significant differences in their shuttling behavior. These results add further support that the orange morphs appear to select different temperatures compared to the white and yellow morphs, and differences in Tpref are not an artifact of differences in movement across the gradient. Taken together, the results of our study suggest that morph thermal preference may be driven, at least in part, by microhabitat differences. Distinct intraspecific morph temperature preferences have myriad implications for how morphs behave and interact (Brock et al., 2022a), as well as their growth rates, physiological performance, reproductive output, and relative morph fitness in different environments (Angilletta et al., 2002; Sinervo & Adolph, 1994; Autumn & DeNardo, 1995; Gray et al., 2008).

We found a significant relationship between body size and Tpref. In our sample, larger lizards had lower preferred body temperatures. Another study on a closely related polymorphic species, *Podarcis gaigeae*, also found a relationship between body size and Tpref, but in the opposite direction: larger lizards preferred higher body temperatures (Runemark et al., 2010; Sagonas et al. 2013b). It is unknown whether or not *P. gaigeae* color morphs differ in body size or thermal preference. Body size may indeed influence lacertid thermoregulation, but not in a clear pattern (Ortega & Martin-Vallejo, 2019). Though Tpref was significantly associated with body size in our study, color morph differences in Tpref remained significant when controlling for body size, suggesting that some other morph-correlated trait, like microhabitat selection, might drive morph differences in Tpref.

Our Tpref results lie within the same thermal range reported in previous studies on *P. erhardii* (Belasen et al., 2017; Pafilis et al., 2019), though orange lizards chose lower Tpref than lizards in these previous studies. Nonetheless, other *Podarcis* species have been shown to select equally low Tpref (Adamopoulou & Valakos, 2005; see Table 2 in Kapsalas et al., 2016). Former investigations on the thermal biology of the Aegean wall lizard treated all individuals indiscriminately in terms of throat color, in contrast to our study here. Our findings demonstrate that color morphs do differ in thermal traits and pave the way for further investigation of the thermoregulation of the species. If color morph is considered, future research may reveal distinct thermoregulatory effectiveness depending on the morph.

In other color polymorphic species, including fish, birds, and lizards, intraspecific color morphs occupy distinct lighting environments to maintain crypsis, avoid predation, and maximize color signaling efficacy (Galeotti et al., 2003; Gray & McKinnon, 2007). In the Indonesian fish *Telmatherina sarasinorum*, male color morphs experience visual environment-contingent sexual selection (Gray et al., 2008). Certain color morphs of *T. sarasinorum* are more conspicuous at different water depths, and thus their space use and frequencies vary across their visually heterogeneous environment (Gray et al., 2008). A comparative analysis of owls, nightjars, and raptors found that color polymorphic species in these groups tend to live in more diverse habitats with open and closed microhabitats compared to monomorphic species (Galeotti & Rubolini, 2004). Environmental resource partitioning, such as occupying different thermal conditions or lighting environments, should reduce intraspecific competition and allow the maintenance of multiple alternative phenotypes within a single population (Forsman & Åberg, 2008). This is common at the interspecific level and evident in other lacertid species that partition their thermal habitat to reduce interspecific competition (Scheers & Van Damme, 2002; Pafilis et al., 2017; Sagonas et al., 2017), but it might also be the case for conspecifics belonging to different color morphs. While morph differences in microhabitat related to lighting environment are well-described, less is known about morph differences in microhabitat use based on thermal conditions. In *P. erhardii*, orange morphs tend to stay close to shady vegetation and use vegetation as refuge more often than white and yellow morphs (Brock & Madden, 2022). Indeed, over several years of fieldwork in the Cyclades islands, we have found brighter orange morphs in more closed, highly vegetated microhabitats (BeVier et al., 2022; Brock et al., 2022b).

If there are constraints on microhabitat availability, variation in color morph preferred body temperature may have consequences on relative color morph fitness, especially for an ectotherm like *P. erhardii*, due to many temperature-dependent biological functions, such as metabolic rate, digestion, growth, and reproduction (Bennett & Dawson, 1976; Pafilis et al., 2007). Lower metabolic rates are associated with lower body temperatures (Bennett & Dawson, 1976). Lower body temperatures in orange lizards may allow them to allocate more ingested energy into growth, which could help explain their larger body size. Furthermore, thermal preferences may affect the geographical distribution of the color morphs. We tend to find few or no orange morphs on the hottest, driest islands with little environmental heterogeneity in the Cyclades (Brock et al., 2022b). If environmental conditions consistently favor a certain morph or morphs over others, non-favored morphs can go extinct, and favored morphs become fixed in the population (West-Eberhard, 1986; Forsman et al., 2008; Corl et al., 2010; Massot et al., 2010). However, the mechanisms by which fixation occurs are not well understood. Across 44 different locations in the Cyclades, *P. erhardii* orange morphs seem to be lost first from 10 islands where only yellow and white morphs remain, one island is fixed for the yellow morph, and 19 islands have white morph fixation (Brock et al., 2022b). Morph loss appears to happen in an ordered fashion - first orange, then yellow, and only white morphs remain with one exception. Parallel ordered morph loss across many *P. erhardii* populations from hot, dry islands could be the result of directional selection that favors morphs who do well in those conditions. Future research in this system should investigate morph thermal physiology and the thermal properties of morph microhabitats to determine the relative availability of those microhabitats across different islands. If orange morphs are tied to cooler, wetter environments, they may not be able to persist where those conditions are unavailable.

## Conclusions

Overall, our results provide more evidence that *P. erhardii* color morphs have alternative trait combinations and for that reason may utilize different environmental resources (BeVier et al., 2022; Brock & Madden, 2022; Brock et al., 2022a). Morph differences in temperature preference have profound implications for morph coexistence and morph life histories, especially in thermally sensitive species like ectotherms. The extent to which different temperature-dependent physiological processes differ between color morphs with different preferred temperatures could have a major influence on how color polymorphism evolves, persists, or erodes. Morph differences in social, sexual, and antipredator behavior may be related to their preferred body temperatures, and future research should investigate these relationships using a combination of observations from natural populations and laboratory studies. The role of thermally heterogeneous environments in the evolution of color morph thermal physiology and the maintenance of color polymorphism should be explored further.

## ACKNOWLEDGEMENTS

We thank Dhruthi Mandavilli, Serey Leakna Nouth, Tanmayi Patharkar, and Jerry Sun for their assistance with catching lizards. We thank the Wang Lab (Anusha Bishop, E. Anne Chambers, Jess McLaughlin, Dan Oliviera, and Erin Westeen) for their helpful feedback on drafts of this manuscript. Thanks to John Adams, whose “*The Dharma at Big Sur*” propelled our writing. A.T. was supported by a travel grant and internship grant from the College of Natural Resources at the University of California, Berkeley. This work was funded by the National Science Foundation PRFB awarded to K.M.B.

